# Strain background, species frequency and environmental conditions are important in determining population dynamics and species co-existence between *Pseudomonas aeruginosa* and *Staphylococcus aureus*

**DOI:** 10.1101/2020.04.21.052670

**Authors:** Selina Niggli, Rolf Kümmerli

**Affiliations:** Department of Quantitative Biomedicine, University of Zürich, Winterthurerstrasse 190, 8057 Zürich, Switzerland

**Keywords:** opportunistic human pathogens, interspecies interactions, polymicrobial infections, ecological factors, community dynamics

## Abstract

Bacterial communities in the environment and in infections are typically diverse, yet we know little about the factors that determine interspecies interactions. Here, we apply concepts from ecological theory to understand how biotic and abiotic factors affect interaction patterns between the two opportunistic human pathogens *Pseudomonas aeruginosa* and *Staphyloccocus aureus*, which often co-occur in polymicrobial infections. Specifically, we conducted a series of short- and long-term competition experiments between *P. aeruginosa* PAO1 (as our reference strain) and three different *S. aureus* strains (Cowan I, 6850, JE2) at three starting frequencies and under three environmental (culturing) conditions. We found that the competitive ability of *P. aeruginosa* strongly depended on the strain background of *S. aureus*, whereby *P. aeruginosa* dominated against Cowan I and 6850, but not against JE2. In the latter case, both species could end up as winners depending on conditions. Specifically, we observed strong frequency-dependent fitness patterns, including positive frequency dependence, where *P. aeruginosa* could dominate JE2 only when common, but not when rare. Finally, changes in environmental (culturing) conditions fundamentally altered the competitive balance between the two species, in a way that *P. aeruginosa* dominance increased when moving from shaken to static environments. Altogether, our results highlight that ecological details can have profound effects on the competitive dynamics between co-infecting pathogens, and determine whether two species can co-exist or invade each others’ populations from rare. Moreover, our findings might parallel certain dynamics observed in chronic polymicrobial infections.

**Importance:** Bacterial infections are frequently caused by more than one species and such polymicrobial infections are often considered more virulent and more difficult to treat than the respective monospecies infections. *Pseudomonas aeruginosa* and *Staphyloccocus aureus* are among the most important pathogens in polymicrobial infections and their co-occurrence is linked to worse disease outcome. There is great interest in understanding how these two species interact with each other and what the consequences for the host are. While previous studies have mainly looked at molecular mechanisms implicated in interactions between *P. aeruginosa* and *S. aureus*, here we show that ecological factors such as strain background, species frequency and environmental conditions are important elements determining population dynamics and species co-existence patterns. We propose that the uncovered principles may also play major roles in infections, and therefore proclaim that an integrative approach combining molecular and ecological aspects is required to fully understand polymicrobial infections.

## Introduction

Bacteria typically live in complex multi-species communities in the environment and associated with host organisms (1–3). The same holds true in the case of disease, as it is increasingly recognized that a majority of bacterial infections are polymicrobial, meaning that they are caused by more than one bacterial species (4, 5). There is great interest in understanding how bacteria interact and how interactions affect a community and the associated hosts (6–8). At the mechanistic level, a multitude of ways have been unraveled through which bacterial species can interact, with mechanisms including cross-feeding, quorum sensing-based signaling, toxin-mediated interference and physical interactions via contact-dependent systems (e.g. type VI secretion system) (9– 11). In the context of disease, a key question is how interactions affect species successions in chronic infections and whether multispecies infections are more virulent and more difficult to treat than the respective monospecies infections, as it is commonly assumed (5, 12–14).

### Studying interactions between the two opportunistic human pathogens

*Pseudomonas aeruginosa* (PA) and *Staphylococcus aureus* (SA) has emerged as a popular and relevant model system (15–17). The reason for this is that the two species often co-occur in infections, including cystic fibrosis (CF) lung and wound infections (18–20). Results from laboratory experiments suggest that PA is the superior species, suppressing growth of SA (21–23) and indeed, PA seems to be a well-equipped competitor. For example, it has been shown that 4-hydroxy-2-heptylquinoline N-oxide (HQNO) released by PA inhibits the electron transport chain of SA and induces the formation of small colony variants (SCVs) (22, 24). Furthermore, the PA endopeptidase LasA is capable of lysing SA cells, a process that releases iron into the environment, potentially providing a direct benefit to PA (21, 25). While it was observed that experimental co-infections of PA and SA seem to be more virulent than the respective monospecies infections (26, 27), evolutionary studies revealed that PA can adapt to the presence of SA (28) and become more benign in the context of chronic co-infections (29, 30). Moreover, of clinical relevance is the observation that PA and SA exhibited increased antibiotic resistance or tolerance when co-cultured compared to being cultured alone (31–33).

In this study, we follow a complementary approach to examine how biotic and abiotic ecological factors influence interactions between PA and SA. Previous work has primarily focused on the molecular mechanisms driving interactions between specific strain pairs under defined laboratory conditions. Here we hypothesize that not only molecular mechanisms, but also ecological factors will have a major impact on species interactions, particularly on community composition and temporal dynamics between species. To test our predictions, we used PA strain PAO1 as our focal strain and asked how (competitive) interactions with SA vary when manipulating: (1) the genetic background of SA; (2) the frequency of SA in competition with PA; and (3) environmental (culturing) conditions.

To vary the genetic background of SA, we competed PA against the three different SA strains Cowan I, 6850 and JE2. These strains fundamentally differ in several characteristics (Table 1). Cowan I is a methicillin-sensitive SA strain (MSSA), which is highly invasive towards host cells, non-cytotoxic and defective in the accessory gene regulator (agr) quorum-sensing system (34). 6850 is another MSSA strain, which is highly invasive, cytotoxic and haemolytic (35–37). Finally, JE2 is a methicillin-resistant (MRSA) USA300 strain, which is highly virulent, cytotoxic and hemolytic (38, 39). Given the tremendous differences between these SA strains, we expect PA performance in competition with SA to vary substantially.

**Table 1.** PA and SA strains used for this study.

| Strain name | Origin | Description | Reference |
| --- | --- | --- | --- |
| ***Pseudomonas aeruginosa***<br>(PA) |  |  |  |
| PAO1::gfp | Wound | Constitutive GFP expression from the chromosome (attTn7::Ptac-GFP). | Our laboratory |
| ***Staphylococcus aureus***<br>(SA) |  |  |  |
| Cowan I | Septic arthritis | MSSA isolate. Highly invasive, but not cytotoxic. Agr-defective. | ATCC 12598 |
| 6850 | Osteomyelitis | MSSA isolate. Highly invasive, cytotoxic and hemolytic. | ATCC 53657 |
| JE2 | Skin and soft tissue infection | USA300 CA-MRSA isolate. Highly virulent, cytotoxic and hemolytic. | NARSA |

CA-MRSA: Community-acquired methicillin-resistant *S. aureus*
MSSA: Methicillin-sensitive *S. aureus*
Agr: Accessory gene regulator

To manipulate strain frequency, we competed PA against SA at three different starting frequencies (1:9 ; 1:1 ; 9:1). Frequency-dependent fitness effects occur in many microbiological systems (40–43). A common pattern is that species have a relative fitness advantage when rare in a population, but not when common (so-called negative frequency dependence). This phenomenon can lead to stable co-existence of competitors. On the other hand, the fitness of a species can also be positive frequency dependent, which means that a species is dominant when common in the community, but not when rare. An important consequence of this pattern is that initially rare species cannot invade an established population.

To manipulate environmental conditions, we changed simple parameters of our culturing conditions. First, we compared the performance of PA against SA strains in shaken liquid vs. viscous medium. Increased environmental viscosity has been shown to increase spatial structure, thereby decreasing strain interaction rates (44–46). Second, we compared the performance of PA against SA in shaken vs. static environments. While static conditions also reduce strain mixing, it further leads to a more heterogeneous environment characterized by gradients from the aerated air-liquid interface down to the microoxic bottom of a culture (47, 48).

In a first set of experiments, we assessed the growth performance of all strains in monoculture under the three different environmental culturing conditions used. Basic growth differences between strains could induce frequency shifts in co-cultures even in the absence of direct interactions. We then performed high-throughput 24 hours batch culture competition experiments between PA and SA using a full-factorial design. All three species combinations were competed at all three starting frequencies under all three environmental conditions (see Figure 1 for an illustration of the workflow). Finally, we followed the temporal dynamics between PA and SA over five days to assess whether results from 24 hours competitions are predictive for more long-term dynamics between species and whether species coexistence is possible. (A copy of this manuscript has been posted on bioRxiv doi: https://doi.org/10.1101/2020.04.21.052670)

**Figure 1.**
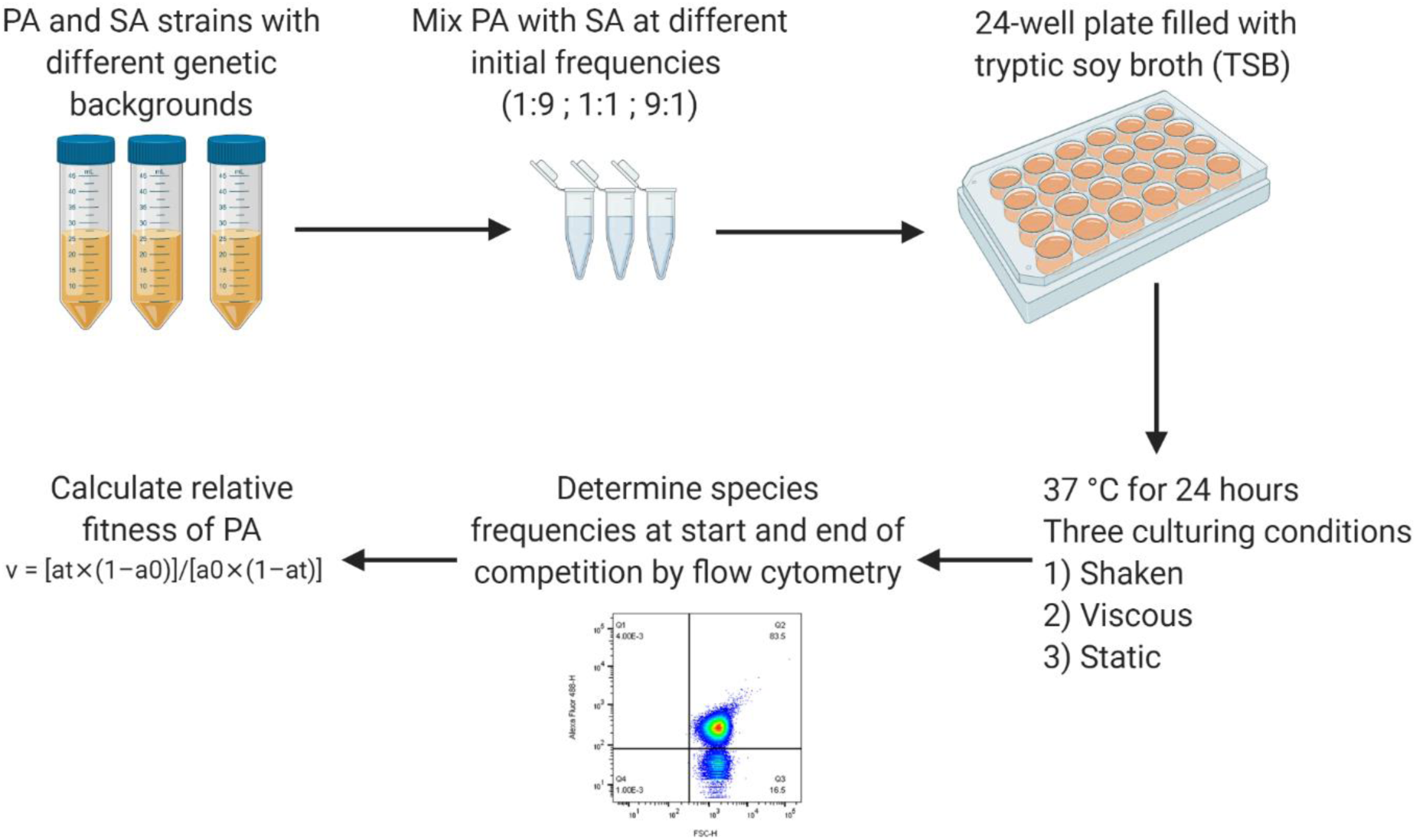
Workflow for the competition experiments. Bacterial overnight cultures were grown in 10 ml TSB in 50 ml falcon tubes for ∼ 16 hours at 37 °C and 220 rpm with aeration. After washing and adjustment of OD_600_ to obtain similar cell numbers for all strains, strain pairs (PA-Cowan I ; PA-6850 ; PA-JE2) were mixed at three different volumetric starting frequencies (1:9 ; 1:1 ; 9:1). Flow cytometry was used to measure the actual starting frequencies. Competitions were started with diluted cultures (OD_600_ = 10^−5^) in 24-well plates filled with 1.5 ml TSB per well. Plates were incubated for 24 hours at 37 °C under three different culturing conditions: shaken (170 rpm), viscous (170 rpm + 0.2% agar in TSB) and static. After the 24 hours competition period, final strain frequencies were measured for each replicate by flow cytometry. Using the initial and final strain frequencies, the relative fitness (v) of the focal strain PA was calculated as v = [a_t_ × (1−a_0_)]/ [a_0_ × (1−a_t_)], where a_0_ and a_t_ are the initial and final frequencies of PA, respectively.

## Results

### PA grows better than SA in monoculture

We used tryptic soy broth (TSB) as the standard medium for all our assays. In this medium, we found that the number of doublings varied significantly among strains during a 24 hours growth cycle under all conditions tested (ANOVA, shaken: F_3,20_ = 10.71, P = 0.0002; viscous: F_3,20_ = 4.12, P = 0.0199; static: F_3,20_ = 20.75, P < 0.0001, Figure 2 and see Table S1 for the full statistical analysis). Under shaken conditions, PA had the highest number of doublings (20.6 ± 0.63, mean ± SD), followed by SA strains 6850 (18.9 ± 1.16), Cowan I (18.1 ± 0.41) and JE2 (18.0 ± 1.18). While PA grew significantly better than all SA strains, the number of doublings did not differ between the three SA strains (TukeyHSD pairwise comparisons: Cowan I vs. 6850, P_adj_ = 0.3673; 6850 vs. JE2, P_adj_ = 0.3096; Cowan I vs. JE2, P_adj_ = 0.9994). Due to its moderate growth advantage, PA is expected to slightly increase in frequency in competition with SA strains under shaken conditions, even in the absence of any direct species interactions.

**Figure 2.**
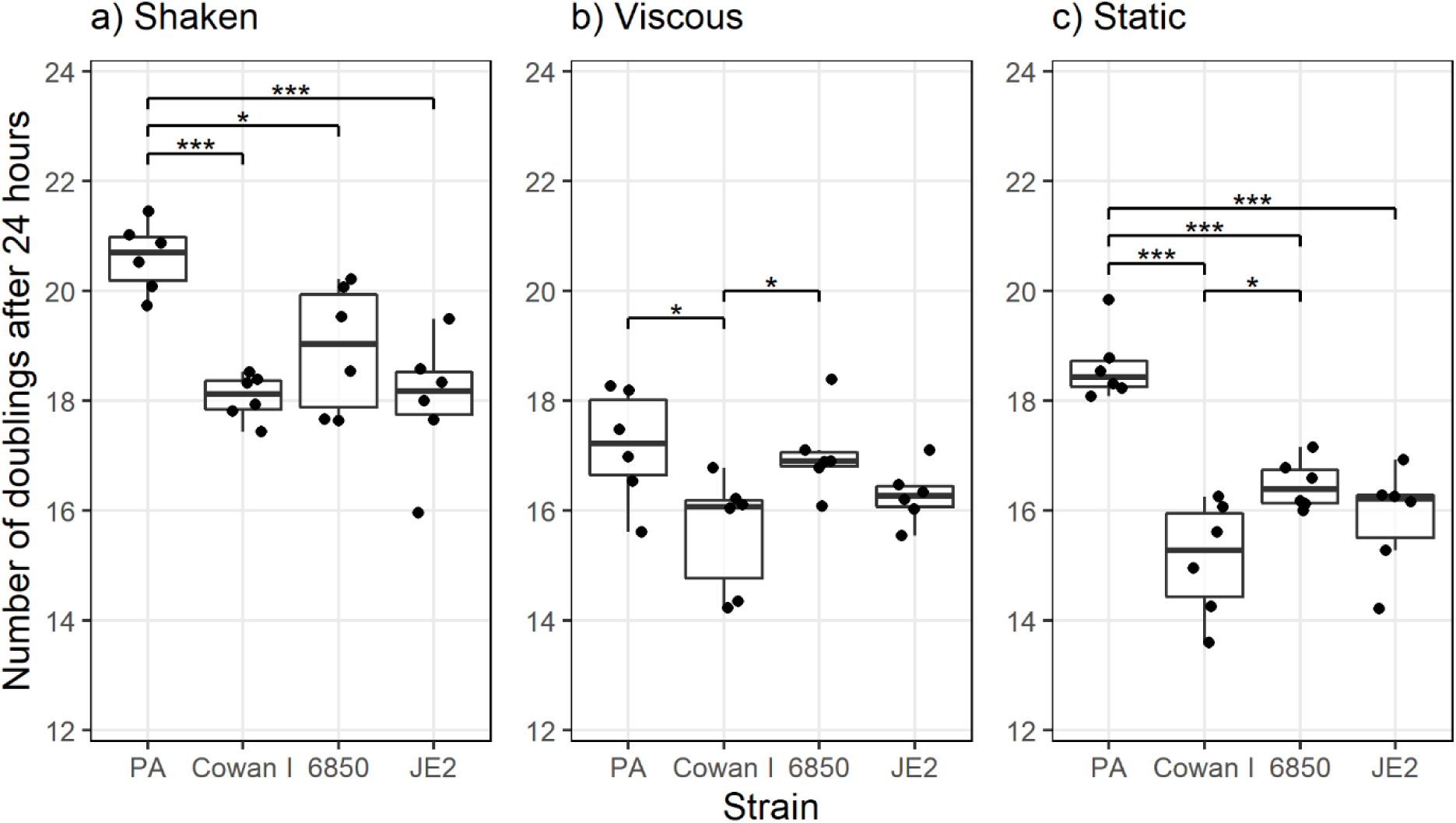
The number of doublings in monoculture is higher for *P. aeruginosa* PAO1 (PA) than for the three *S. aureus* strains (Cowan I, 6850, JE2) under most conditions. Strains were grown as monocultures in TSB for 24 hours at 37 °C under the same conditions and using the same starting OD_600_ as for the competition experiments. The box plots show the median (bold line) with the first and the third quartiles. The whiskers cover the 1.5* inter-quartile range (IQR) or extend from the lowest to the highest value if they fall within the 1.5* IQR. Data are shown from two independent experiments with three replicates each. * p < 0.05, *** p < 0.001, pairwise comparisons without bars are all not significant.

Under viscous conditions, PA had the highest number of doublings as well (17.2 ± 1.02), followed by 6850 (17.0 ± 0.76), JE2 (16.3 ± 0.52) and Cowan I (15.6 ± 1.06). However, differences in the number of doublings were only significant between PA and Cowan I and between Cowan I and 6850 (TukeyHSD pairwise comparisons: PA vs. Cowan I, P_adj_ = 0.0265; Cowan I vs. 6850, P_adj_ = 0.0496). Thus, based on growth rate differences alone, one would expect PA to increase in frequency in competition with Cowan I but not in competition with the other two SA strains.

Under static conditions, PA again showed the highest number of doublings (18.6 ± 0.64) followed by 6850 (16.5 ± 0.45), JE2 (15.9 ± 0.96) and Cowan I (15.1 ± 1.05). PA grew significantly faster than all the three SA strains (TukeyHSD pairwise comparisons: PA vs. Cowan I, P_adj_ < 0.0001; PA vs. 6850, P_adj_ = 0.0009; PA vs. JE2, P_adj_ < 0.0001), and one would therefore expect PA to substantially increase in frequency against all three SA strains under static conditions.

### Genetic background, strain frequency and environmental factors all influence competition outcomes

The full-factorial design allowed us to simultaneously analyze the impact of SA strain genetic background, starting frequency, and culturing condition on the competitive outcomes between PA and SA strains. Our linear statistical model yielded a significant triple interaction between the three manipulated factors (strain genetic background, starting frequency and culturing condition; ANCOVA: F_4,509_ = 3.41, P = 0.0091). While this shows that all three manipulated factors influence the competitive outcomes between PA and SA in complex ways, the triple interaction makes it difficult to tease apart the various effects. The statistical procedure for such cases is to split the model into sub-models. We followed this approach by first analyzing separate models for each of the three environmental conditions (shaken, viscous, static), and then split models according to SA strain background to test for differences between environmental conditions.

### The competitive ability of PA depends on the SA strain genetic background

Under all three environmental conditions, we found that the relative fitness of PA significantly depended on the SA strain background (ANCOVA, shaken: F_2,170_ = 90.87, P < 0.0001; viscous: F_2,168_ = 116.76, P < 0.0001; static: F_2,170_ = 56.52, P < 0.0001; Figure 3). Against Cowan I (Figure 3, column 1), we noted that PA consistently won the competitions across all starting frequencies and culturing conditions. SA strain 6850 (Figure 3, column 2) turned out to be more competitive than Cowan I under shaken conditions (t_176_ = -6.74, P < 0.0001), while it lost similarly against PA under viscous and static conditions (viscous: t_174_ = 0.78, P = 0.4350; static: t_176_ = -1.99, P = 0.0482). In contrast, JE2 was the most competitive SA strain in our panel (Figure 3, column 3), performing significantly better than the other two SA strains under all conditions (see Table S2 for the full statistical analysis), and outcompeted PA under shaken and viscous conditions.

**Figure 3.**
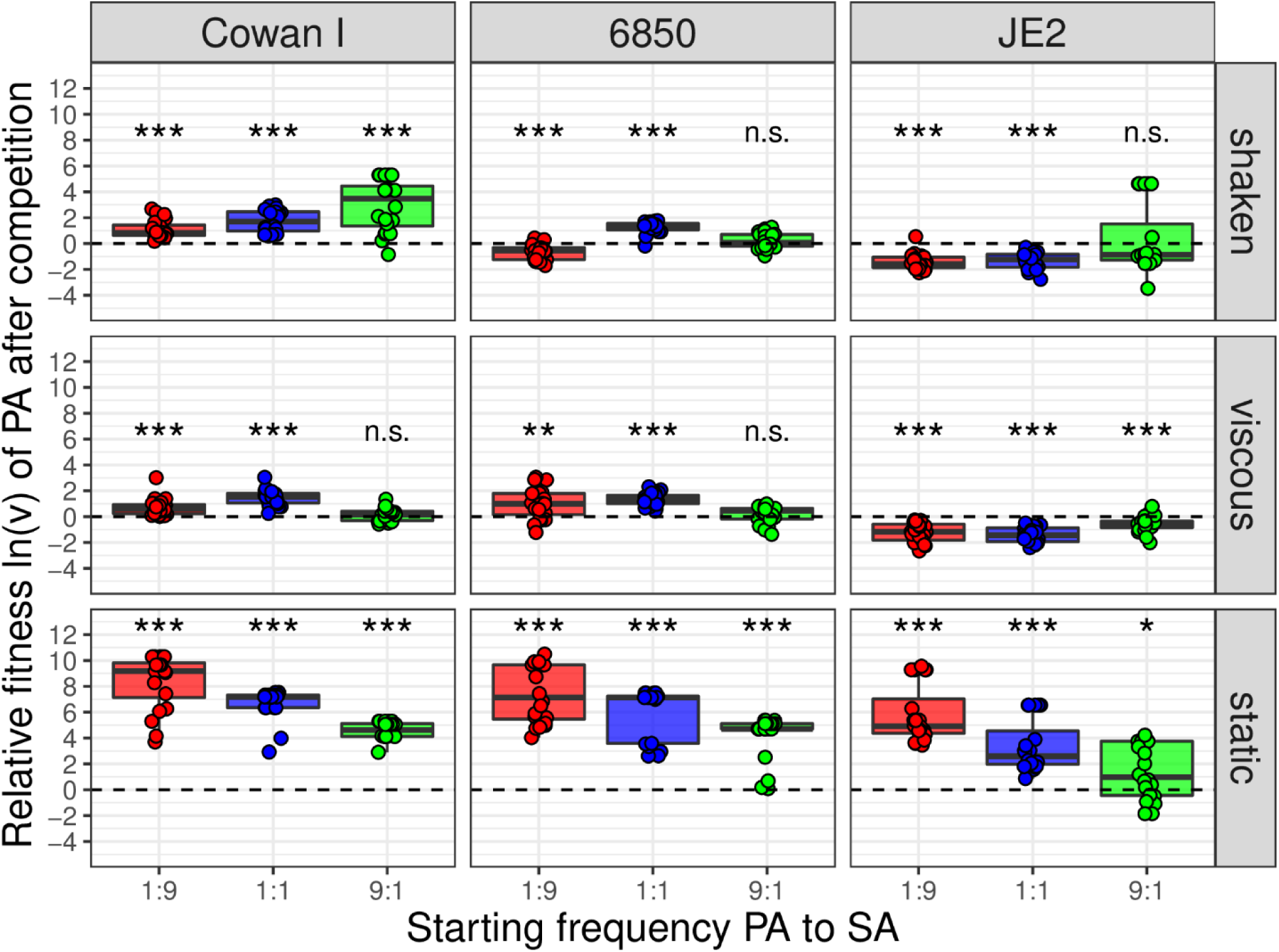
Relative fitness ln(v) of *P. aeruginosa* PAO1 (PA) after 24-hours competitions against three different *S. aureus* (SA) strains (Cowan I, 6850, JE2) at three different starting frequencies (1:9 ; 1:1 ; 9:1) and across three different environmental conditions (shaken, viscous, static). Values of ln(v) < 0, ln(v) > 0, or ln(v) = 0 (dotted line), indicate whether PA lost, won, or performed equally well in competition against the respective SA strain. The box plots show the median (bold line) with the first and third quartiles. The whiskers cover the 1.5* inter-quartile range (IQR) or extend from the lowest to the highest value if they fall within the 1.5* IQR. Each strain pair/culturing condition/starting frequency combination was repeated 20 times (four experiments featuring five replicates each). Asterisks indicate whether the relative fitness of PA is significantly different from zero in a specific treatment (one-sample t-tests with p-values corrected for multiple comparisons using the false discovery rate method: n.s. = not significant, * p < 0.05, ** p < 0.01, *** p < 0.001). Detailed information on all statistical comparisons are provided in Table S3.

### The competitive ability of PA depends on its starting frequency in the population

We found that the starting frequency of the two competitors had varying but always significant effects on the competitive ability of PA (ANCOVA, shaken: F_1,170_ = 52.81, P < 0.0001; viscous, interaction with strain background: F_2,168_ = 10.05, P < 0.0001; static: F_1,170_ = 162.32, P < 0.0001). Under shaken conditions (Figure 3, row 1), we observed that the relative fitness of PA increased when initially more common, thus following a positive frequency-dependent pattern. Under viscous conditions (Figure 3, row 2), the same positive frequency-dependent effect was only observed when PA competed with JE2. In competition with Cowan I or 6850, we noted that the relative fitness of PA peaked at intermediate starting frequencies. Under static conditions (Figure 3, row 3), we observed a pattern opposite to the one seen under shaken conditions for all strain pair combinations. The relative fitness of PA decreased when initially more common, thus following a negative frequency-dependent pattern (see Table S2 for the full statistical analysis).

### The competitive ability of PA is highest under static conditions

Next, we compared the competitive outcomes among the different culturing conditions (shaken, viscous and static) for each strain combination separately. For all strain combinations, the culturing condition significantly affected competition outcomes (ANCOVA, Cowan I: F_2,168_ = 461.73, P < 0.0001; 6850: F_2,167_ = 428.16, P < 0.0001; JE2: F_2,168_ = 199.95, P < 0.0001). In competition with all three SA strains, we found that the relative fitness of PA was significantly higher under static compared to shaken conditions (Cowan I: t_174_ = 19.99, P < 0.0001; 6850: t_174_ = 17.99, P < 0.0001; JE2: t_174_ = 15.39, P < 0.0001). In contrast, there were no significant differences in the relative fitness of PA between shaken and viscous conditions for Cowan I (t_174_ = 0.91, P = 0.3644) and JE2 (t_174_ = 0.82, P = 0.4117), while against 6850, PA was more competitive under viscous than shaken conditions (t_174_ = 3.53, P = 0.0005) (see Table S2 for the full statistical analysis).

### Temporal dynamics between PA and SA

In a next experiment, we competed PA and SA strains over five days under shaken conditions using the same three starting frequencies and by transferring cultures to fresh medium every 24 hours. The aim of this experiment was to follow the more long-term species dynamics and to assess whether stable coexistence between PA and SA can arise.

In competition with Cowan I, we found PA to be the dominant species (Figure 4a). It strongly increased in frequency already at day 1 under all starting frequencies and almost completely outcompeted Cowan I by day 3 (i.e., Cowan I remained below detection limit). Thus, we could not observe coexistence between PA and Cowan I. In competition with 6850, we observed similar population dynamics (Figure 4b). PA strongly increased in frequency from day 1 onwards at all starting frequencies and after three days, the bacterial populations almost entirely consisted of PA. Only in 10 out of 30 populations, 6850 managed to persist at very low frequencies by day 5 (< 3% in nine cases, and 13% in one case). In competition with JE2, we found community trajectories that were strikingly different from the other two strain combinations (Figure 4c). First, we observed that JE2 was a strong competitor, keeping PA at bay in many populations during the first 24 hours of the experiment. Following day 1, community dynamics followed positive frequency-dependent patterns. In all populations with intermediate or high PA starting frequencies, PA became the dominant species, and SA was recovered at low frequency in only a minority of populations by day 5 (3 out of 20 at < 10% of the population). In stark contrast, in populations where PA was initially rare, it did not increase in frequency, could not invade the SA populations and remained at a low frequency (< 10%) throughout the 5 days.

**Figure 4.**
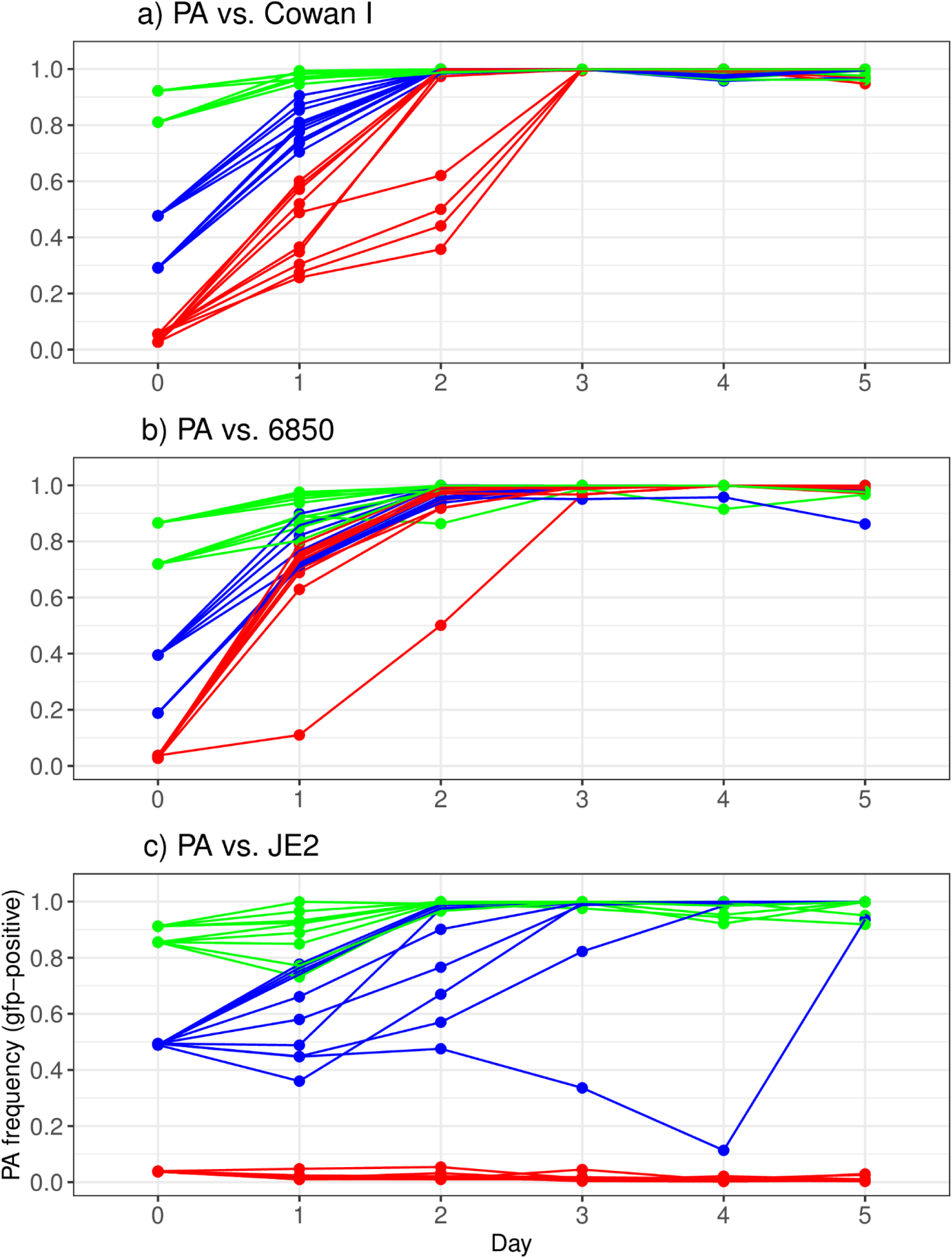
Multi-day competitive dynamics between *P. aeruginosa* PAO1 (PA) and the three *S. aureus* strains (a) Cowan I, (b) 6850 and (c) JE2 under shaken conditions. Competitions started at three volumetric starting frequencies of PA:SA (red 1:9, blue 1:1, green 9:1). Community composition was followed over five days with daily transfer of diluted cultures to fresh TSB medium. Strain frequencies were assessed using flow cytometry. The experiment was carried out two times with five replicates per treatment combination and experiment.

## Discussion

*Pseudomonas aeruginosa* (PA) and *Staphylococcus aureus* (SA) frequently occur together in polymicrobial infections, where they cause severe host damage and lead to increased morbidity and mortality in patients (14, 49, 50). Consequently, there is high interest in understanding how PA and SA interact and how their interactions may influence disease outcome (12, 15). While most previous studies have focused on molecular aspects (51, 52), we here examined how a set of ecological factors affect competitive interactions between the two species. Our study, carried out in an *in vitro* batch culture system, revealed that: (i) the competitive ability of PA varied extensively as a function of the genetic background of SA; (ii) there were strong frequency-dependent fitness patterns, including positive-frequency dependent relationships where PA could only dominate a particular SA strain (JE2) when common, but not when rare; and (iii) changes in environmental (culturing) conditions fundamentally affected the competitive balance between the two species. The key conclusion from our results is that ecology matters, and that variation in biotic and abiotic factors affect interactions between pathogenic bacterial species. This is most likely not only the case in *in vitro* systems, but also in the context of polymicrobial infections.

PA has often been described as the dominant pathogen possibly displacing SA in infections (21, 53–55). Our results support this view, as PA dominated over SA under many conditions in 24 hours and 5-day competition experiments (Figures 3 and 4). However, PA did not always emerge as the winner and its success significantly varied in response to the genetic background of SA. Specifically, JE2 was the strongest competitor, followed by 6850 and Cowan I. While differences in monoculture growth performance can explain why PA dominates in many cases, they cannot explain the variation in competitive abilities among SA strains, because the three SA strains grew similarly under all conditions (Figure 2). PA vs. JE2 makes the strongest case for a mismatch between growth performance in mono-vs. mixed cultures, as PA grew significantly better than JE2 in monoculture under shaken conditions, but typically lost the competition in mixed cultures in this environment (Figures 2 and 3). This suggests that apart from resource competition via growth rate differences, other factors must contribute to the competitive ability of SA strains towards PA. Such factors could include interference mechanisms. For example, it is known that genetically different SA strains widely differ in virulence among each other (56–58). Interestingly, we found that the competitive ability of our three SA strains against PA correlated with their reported virulence level in infections (34, 36, 39). This could indicate that factors important for SA virulence (e.g. toxins or secreted enzymes) might also be involved in interactions with competitor bacteria. For JE2 and related USA300 isolates, there are many genetic determinants known to be important for their success as opportunistic human pathogens (59). Among them are the cytotoxin Panton-Valentine leukocidin (PVL), the arginine catabolic mobile element (ACME) and the phenol soluble modulins (PSMs) (60). Derivatives of PSMs have previously been shown to exhibit inhibitory activity against *Streptococcus pyogenes* (61). The authors of this work suggested that high production of PSMs might not only benefit SA in host colonization, but also in competition against co-infecting pathogens. Thus, it seems plausible that the USA300 derivative JE2 deploys a similar mechanism against PA in our competition experiments. Strain 6850 showed intermediate competitiveness against PA. As Cowan I, 6850 is a MSSA strain, but it is known to be more virulent than Cowan I and therefore likely produces certain substances that could also be important in competition with PA (34, 36). Conversely, Cowan I is known to have a nonfunctional accessory gene regulator (agr) quorum sensing system (34). The agr controls most virulence determinants in SA (62). If virulence determinants also played a role in interspecies competition, then this could explain why Cowan I turned out to be the least competitive SA strain against PA.

Important to note is that we only manipulated the SA but not the PA strain background, so that we cannot draw conclusions on genotype-by-genotype interactions. Such interactions are likely to play a role as evidenced by previous studies showing that genetically diverse PA clinical isolates widely differ in their ability to inhibit SA (29, 63, 64). More recently, it has also been shown that SA clinical isolates vary in their interactions with PA, from being highly sensitive to completely tolerant against PA-mediated effects (65). One aim of our future work is to follow up these proposed mechanistic leads and test some of the outlined hypotheses above in order to explain differences in the competitive abilities between SA strains towards PA.

Another insight from our experiments is that the competitive ability of PA often depended on its starting frequency in the population, and that the type of frequency-dependent interactions (positive or negative) varied across environmental conditions (Figure 3). Our purpose was to compare three experimentally defined starting frequencies to mimic what is happening when a species is either rare (1:9), at parity with its competitor (1:1) or dominant (9:1). Under natural conditions, including infections, species frequencies could of course be more extreme, and invasion from rare could start at species frequencies that would be below the detection limit of our methods. In our experiments, we observed that under static conditions, the relative fitness of PA declined when more common in the population, but PA still won at all frequencies. This pattern is common for a highly dominant species that drives a competitor to extinction (66). Its decline in relative fitness simply reflects the fact that the room for further absolute frequency gains is reduced when a high frequency is already reached. In stark contrast, under shaken conditions, we found that the relative fitness of PA increased when it was more common in the population. Against Cowan I and 6850, this positive frequency-dependent fitness pattern did not affect the long-term community dynamics and PA won at all frequencies (Figure 4a+b). Against JE2, however, the 24 hours competition data suggest that, in most cases, PA cannot invade populations when initially rare and this is exactly what we observed in the long-term experiments: when its initial frequency was below 10%, PA did not increase in frequency, while it always fixed in the population or reached very high frequencies when initially occurring above 10%. There were two additional interesting observations with regards to PA-JE2 long-term dynamics. First, there were no major changes in PA frequency relative to JE2 during the first 24 hours (compatible with the competition assay data in Figure 3), and clear positive-frequency dependent patterns only emerged from 48 hours onwards. One possible explanation for this pattern is that PA is initially naïve, but then senses and mounts a more competitive response over time (67). Similarly, SA might also respond, for example through increased formation of small colony variants (SCVs), which are known to be induced by inhibitory exoproducts released by PA (22, 24). Second, one replicate (starting frequency 1:1) did not follow the above rules: PA continuously dropped in frequency until day 4 (11%) and then sharply increased to 93% on day 5. This frequency “zigzag” pattern is an indicator of antagonistic co-evolution (68), where the spread of a beneficial mutation in one species (SA) is followed by a counter-adaptation in the competing species (PA). It therefore seems that such evolutionary dynamics can already occur within relatively short periods of time. This finding supports evidence from studies on clinical PA isolates, which showed patterns of adaptations towards SA favoring co-existence between the two species over time (29, 30).

Our results further show that the competitive ability of PA is profoundly influenced by environmental (culturing) conditions (Figure 3). The largest differences arose between shaken and static culturing conditions with PA being most competitive in the latter environment. PA is known to be metabolically versatile, it is motile and grows well under microoxic conditions (69, 70). Static conditions introduce strong oxygen and nutrient gradients, and our results from monoculture growth show that PA grows better under these conditions than SA (Figure 2). This is certainly part of the reason why PA ends up as the competition winner against all SA strains under static conditions. With regard to medium viscosity, we initially hypothesized that increased spatial structure could temper competitive interactions and favor species co-existence, as competitors are spatially more segregated from each other (44, 71, 72). However, we found no support for this hypothesis as the competitive ability of PA did not much differ between shaken and viscous environments. While the spatial structure, introduced through the addition of agar to the liquid growth medium, had significant effects on within-species social interactions in other study systems (66, 73), it did not appreciably affect the between-species interactions in our setup. One reason might be that the degree of spatial structure introduced (0.2% agar in TSB) was simply not high enough to see an effect. This could especially be true if toxins were involved in mediating interactions – small molecules that can freely diffuse and target competitors that are not physically close-by.

We argue that our results, even though they stem from an *in vitro* system, could have at least three important implications for our understanding of polymicrobial infections. First, we show that the biological details of the strain background matter and determine who is dominant in a co-infection and whether co-existence between species is possible. Thus, we need to be careful not to overinterpret interaction data from a single PA-SA strain pair and conclude that the specific details found apply to PA-SA interactions in general. Second, there might be strong order effects, such that the species that infects a host first cannot be invaded by a later arriving species. This scenario applied to the interactions between PA and SA strain JE2, which were both unable to invade populations of the other species from rare. Finally, local physiological conditions at the infection site, like the degree of spatial structure or oxygen supply, can shift the competitive balance between species. This suggests that infections at certain sites might be more prone than others to polymicrobial infections, or to experience ecological shifts from one pathogen to another. To sum up, we wish to reiterate our take home message that the ecology of interactions between pathogens should receive more attention and may explain so far unresolved aspects of polymicrobial infections.

## Materials and Methods

### Bacterial strains, media and growth conditions

We used the *Pseudomonas aeruginosa* (PA) strain PAO1 (74) as our PA reference strain and the *Staphylococcus aureus* (SA) strains Cowan I, 6850 and JE2 for all experiments (Table 1). To distinguish PA from SA strains, we used a variant of our PA strain PAO1, which constitutively expresses the green fluorescent protein, from a single-copy gene (attTn7::ptac-gfp), stably integrated in the chromosome (75, 76). We chose the rich laboratory medium tryptic soy broth (TSB, Becton Dickinson) for all our experiments, because it supports growth of all the strains used. Bacterial stocks were prepared by mixing 50% of culture with 50% of a 85% glycerol solution and were stored at -80 °C. For all experiments, overnight cultures were grown in 10 ml TSB in 50 ml falcon tubes for ± 16 hours at 37 °C and 220 rpm with aeration. After centrifugation and removal of the supernatant, we washed bacterial cells using 10 ml 0.8% NaCl solution and adjusted the OD_600_ (optical density at 600 nm) to obtain similar cell numbers per ml for each strain. All media, buffer and washing solutions were sterilized by autoclaving at 121°C for 20 minutes and subsequently stored at room temperature in the dark. For all experiments, blanks were used to ensure the sterility of the media during experimentation.

### Calculating number of doublings for each strain in monoculture

To assess the number of doublings of each strain in monoculture, we grew our strains in TSB (or TSB + 0.2% agar, respectively) under the same conditions and using the same starting OD_600_ as for the competition experiments (see below). We serially diluted cells at the start (t_0_) and after 24 hours (t_24_), and plated aliquots on TSB + 1.2% agar. The plates were incubated overnight at 37 °C and colony-forming units (CFUs) counted for both timepoints on the following day. We estimated the number of doublings (D) for each strain as D = (ln(x_24_/x_0_))/ln(2), where x_0_ and x_24_ are the initial and the final CFU/ml, respectively (23). We performed this experiment two times with three replicates per strain per experiment.

### Competition experiments

To initiate competitions, we mixed PA and SA strain pairs at three different starting frequencies (1:9, 1:1, 9:1) from washed and OD_600_-adjusted overnight cultures (see above). Competitions occurred in 24-well plates filled with 1.5 ml TSB per well. The starting OD_600_ of both mixed and monocultures was 10_-5_. Monocultures of each strain served as controls in each experiment. We incubated plates for 24 hours at 37 °C under three different culturing conditions: shaken (170 rpm), viscous (170 rpm with 0.2% agar in TSB) and static. Prior and after the 24 hours competition period, we estimated the actual strain frequencies for each replicate using flow cytometry. We performed four independent experiments each featuring five replicates for each strain/starting frequency/condition combination. A graphical representation of the competition workflow is provided in Figure 1.

To follow community dynamics over time, we set up competitions in the same way as described above. After the first 24 hours of competition, we diluted cultures 1:10,000 into fresh TSB medium. This process was repeated for five consecutive days. Strain frequencies were assessed using flow cytometry prior and after each 24 hours competition cycle. We carried out two independent experiments for each strain pair and starting frequency combination with 5 replicates per strain pair and frequency.

### Flow cytometry to estimate relative species frequency

We assessed the relative strain frequencies at the beginning and at the end of each competition using a BD LSR II Fortessa flow cytometer (flow cytometry facility, University of Zürich) and the FlowJo™ software (BD, Bioscience) for data analysis. As our PA strain expresses a constitutive gfp tag, PA cells could unambiguously be distinguished from the gfp-negative SA cells with a blue laser line (excitation at 488 nm) and the FITC channel (emission: mirror 505 longpass, filter 530/30) (see supplementary Figure 1). Cytometer Setup and Tracking settings of the instrument were used for each experiment and the threshold of particle detection was set to 200 V (lowest possible value). We diluted cultures appropriately in sterile-filtered 1x phosphate buffered saline (PBS, Gibco, Thermo Fisher) and recorded 100,000 events with a low flow rate. The following controls were used for data acquisition in every experiment: 1) PBS blank samples (to estimate number of background counts of the flow cytometer), 2) untagged monocultures (negative fluorescence control, used to set a fluorescence threshold in FlowJo™) and 3) constitutive gfp-expressing monocultures (positive fluorescence control, set to 100% gfp-positive cells). Using our fluorescence threshold, we extracted the percentage of gfp-positive cells for each sample and scaled these values to the positive fluorescence control. The resulting percentage corresponds to the frequency of PA present in the respective replicate. Initial and final strain frequencies were used to calculate the relative fitness (v) of the focal strain PA as v = [a_t_ × (1−a_0_)]/ [a_0_ × (1−a_t_)], where a_0_ and a_t_ are the initial and final frequencies of PA, respectively (43). We ln-transformed all relative fitness values to obtain normally distributed residuals. Values of ln(v) > 0 or ln(v) < 0 indicate whether the frequency of the focal strain PA increased (i.e. PA won the competition) or decreased (i.e. PA lost competition) relative to its SA competitor.

We know from previous experiments in our laboratory that due to the gfp tag, our PA strain does have a slight fitness defect in competition with its untagged parental strain (ln(v) = −0.358 ± 0.13, mean ± 95% CI, see (66)). As we consistently used the same gfp-tagged PA strain for all experiments in this study, results are fully comparable among treatments.

To test whether flow cytometry counts (measuring all viable and non-viable cells) correlate with CFUs (measuring only viable cells), we serially diluted and plated initial and final strain frequencies from competitions performed under shaken conditions for all three strain combinations on TSB + 1.2% agar. We compared the obtained CFUs with the flow cytometry counts from the same samples and found strong positive correlations for the strain frequency estimates between the two methods (see supplementary figure 2). This means that flow cytometry adequately measures strain frequencies and that the two methods (flow cytometry and CFU counts) yield similar results.

### Statistical analysis

All statistical analyses were performed with R Studio version 3.6.1. We used analysis of variance (ANOVA) and Tukey’s HSD to compare number of doublings in monocultures of PA and SA. To test whether the relative fitness of PA varies in response to the SA strain genetic background, starting frequency and culturing conditions, we first built a factorial analysis of co-variance (ANCOVA), with SA strain genetic background and culturing conditions as factors and the starting frequency as covariate. We further included ‘experimental block’ as an additional factor to account for variation between experiments. This full model yielded a significant triple interaction between SA strain genetic background, starting frequency and culturing condition. We therefore split the full model into a set of ANCOVA sub-models, separated either by culturing condition (shaken, viscous, static) or by SA strain genetic background (Cowan I, 6850, JE2). For post-hoc pairwise comparisons between culturing conditions or SA strains in the sub-models, we removed ‘experimental block’ as additional factor from the model. To test whether PA relative fitness is significantly different from zero under a given strain/starting frequency/condition combination, we performed one sample t-tests and used the false discovery rate method to correct p-values for multiple comparisons (77). To compare strain frequencies obtained by flow cytometry with those obtained by plating (CFUs), we used Pearson correlation analysis. For all data sets, we consulted Q-Q plots and results from the Shapiro-Wilk test to ensure that our residuals were normally distributed. Summary tables for linear models and t-tests used to analyze Figures 2 and 3 can be found in the supplemental material (Tables 1-3).

## Data availability

All raw data sets have been deposited in the figshare repository (DOI will be provided upon the acceptance of the manuscript).

## Acknowledgements

We thank Markus Huemer (University Hospital of Zürich) for providing *S. aureus* strains and the flow cytometry facility (University of Zürich) for technical support and maintenance of resources. This project has received funding from the European Research Council (ERC) under the European Union’s Horizon 2020 research and innovation programme (grant agreement no. 681295) to RK. Illustration for Figure 1 was created using BioRender (www.biorender.com).

## Author contributions

S.N. and R.K. designed research, S.N. performed research, S.N. and R.K. analysed data and wrote the paper.

