## Supplementary figures 1, 2 for "Strain background, species frequency and environmental conditions are important in determining population dynamics and species co-existence between *Pseudomonas aeruginosa* and *Staphylococcus aureus*"

**a) PA strain PAO1 untagged** **b) PAO1::gfp** **c) 40% PAO1::gfp + 60% SA**

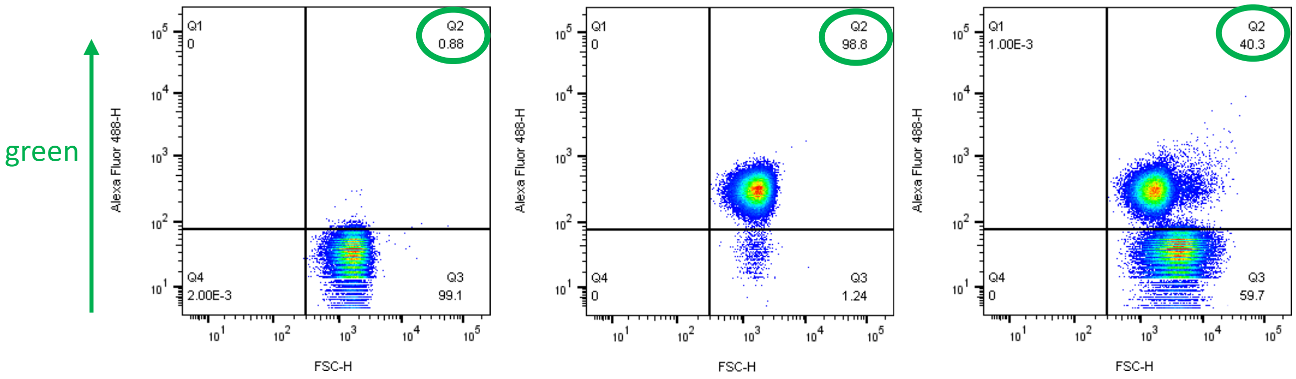


**Supplementary Figure S1.** Flow cytometry plots explaining how we distinguished gfp-positive from gfp-negative (untagged) cells. X-axis shows forward scatter (FSC), a proxy for particle size and y-axis shows Alexa Fluor 488 (gfp) expression. a) PA untagged (negative fluorescence control), used to set a threshold for gfp-positive vs. gfp-negative cells in FlowJo. b) PA tagged chromosomally with gfp, by applying the same fluorescence threshold, shows 98.8% gfp-positive cells. This percentage was set as 100% and all mixed culture samples were scaled to this value. c) Plot showing a mixture of approximately 40% PA + 60% SA cells. Gfp-positive PA cells can clearly be distinguished from gfp-negative SA cells. Fluorescence thresholds were newly determined for every experiment.

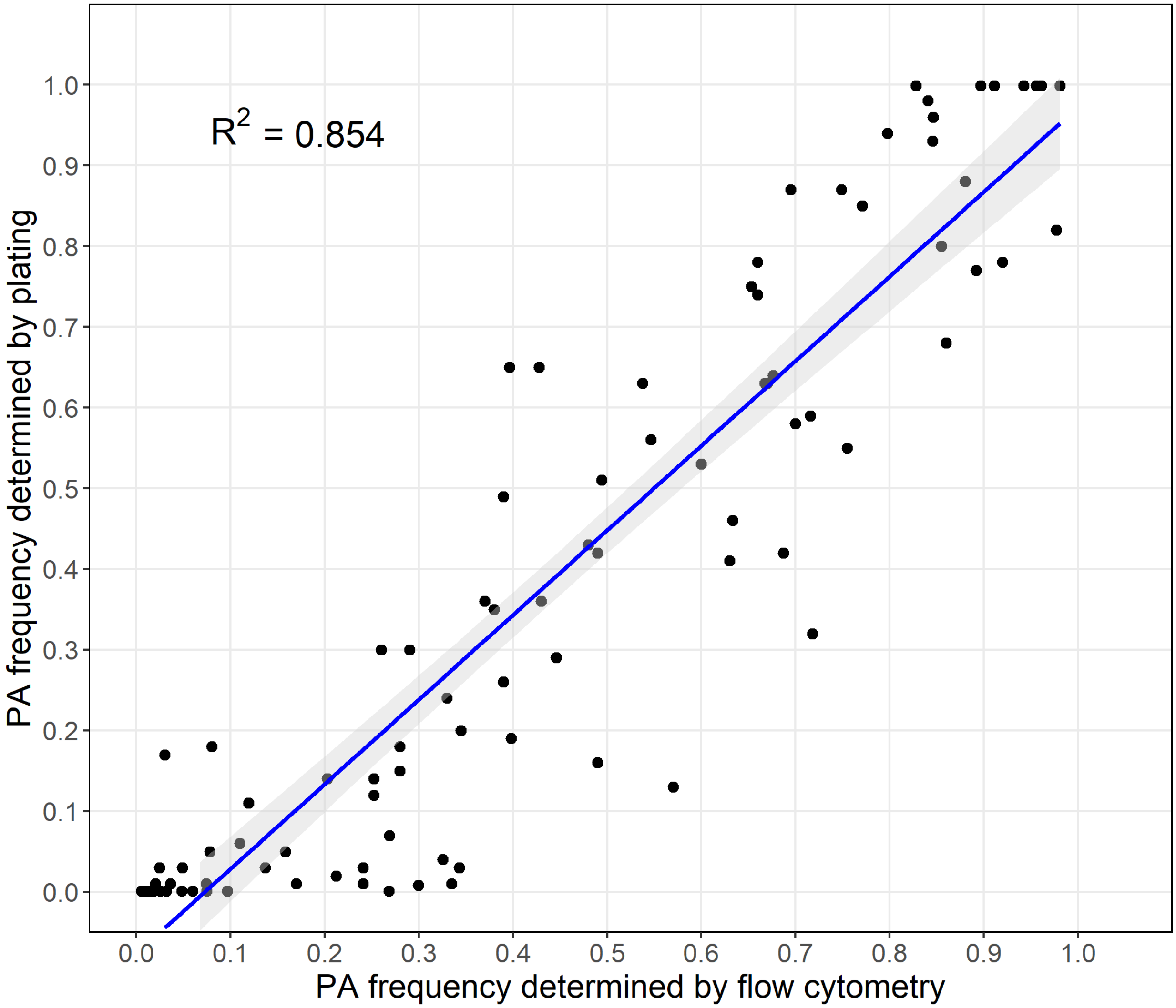


**Supplementary Figure S2.** Correlation between PA frequency determined by flow cytometry (x-axis) and PA frequency determined by plating (y-axis). We did this control experiment for all strain pair combinations and data is from eight independent experiments. Frequencies determined by flow cytometry correlate well with frequencies determined by plating (Pearson correlation coefficient $R^2$ = 0.854).
